# A plasmid-based *Escherichia coli* gene expression system with cell-to-cell variation below the extrinsic noise limit

**DOI:** 10.1101/192963

**Authors:** Zach Hensel

## Abstract

Experiments in synthetic biology and microbiology can benefit from protein expression systems with low cell-to-cell variability (noise) and expression levels precisely tunable across a useful dynamic range. Despite advances in understanding the molecular biology of microbial gene regulation, many experiments employ protein-expression systems exhibiting high noise and nearly all-or-none responses to induction. I present an expression system that incorporates elements known to reduce gene expression noise: negative autoregulation and bicistronic transcription. I show by stochastic simulation that while negative autoregulation can produce a more gradual response to induction, bicistronic expression of a repressor and gene of interest can be necessary to reduce noise below the extrinsic limit. I synthesized a plasmid-based system incorporating these principles and studied its properties in *Escherichia coli* cells, using flow cytometry and fluorescence microscopy to characterize induction dose-response, induction/repression kinetics and gene expression noise. By varying ribosome binding site strengths, expression levels from 55— 10,740 molecules/cell were achieved with noise below the extrinsic limit. Individual strains are inducible across a dynamic range greater than 20-fold. Experimental comparison of different regulatory networks confirmed that bicistronic autoregulation reduces noise, and revealed unexpectedly high noise for a conventional expression system with a constitutively expressed transcriptional repressor. I suggest a hybrid, low-noise expression system to increase the dynamic range.

## Introduction

Experiments in microbiology commonly call for recombinant expression of a protein of interest with expression levels typical of endogenous proteins (~10—10,000 molecules/cell)^1^. However, many experiments today utilize expression systems that are best suited for one-time induction of protein overexpression—systems that exhibit nearly all-or-none response to induction and, often, interference with cellular metabolic networks. For example, consider one single-molecule experiment requiring the expression of two different fluorescent proteins, each on the order of 100 molecules per *Escherichia coli* cell^2^. The complicated growth protocol required optimizing media choices, inducer concentration, washing steps, and induction times. Yet, in the subsequent experiment, only a small fraction of cells contained a useful amount of both proteins of interest. In this case and many like it, a protein expression system with low noise at low expression levels would reduce the time needed to optimize sample preparation protocols and increase data throughput. A low-noise expression system could also improve yields in applications where protein aggregation is a challenge^3^ by making it easier to tune expression so most cells are producing as much protein as possible without being so high as to trigger aggregation; some commonly used induction systems exhibit large cell-to-cell variation even at high expression levels when analyzed at the single-cell level^4^.

Autoregulation is a common motif in prokaryotic gene regulatory networks^5^. Steady-state fluorescence experiments have shown that negative autoregulation by the tetracycline-inducible transcriptional repressor TetR-EGFP can reduce gene expression noise in *Escherichia coli*^6,7^, which was suggested to result from dosage compensation for plasmid copy number^8^. A similar network was implemented in *Saccharomyces cerevisiae* and compared to regulation of EGFP by constitutively expressed TetR; negative autoregulation reduced expression noise and linearized the inducer dose-response^9^. With some caveats, noise can be decomposed to the sum of factors “intrinsic” to expression of the gene of interest (arising from the randomness of binding, transcription, translation, and degradation) and “extrinsic” cell-wide properties (*e.g.* gene dosage and global transcription/translation rates)^8^. Timelapse experiments showed that negative autoregulation can counter long-lived extrinsic noise^10^. Negative autoregulation has also been shown to shift noise in gene expression to higher frequencies^11^ (*i.e.* an autoregulated gene can respond more quickly to fluctuations, and downstream processes that respond to the integrated signal of the autoregulated gene over time can exhibit less noise).

In cases where autoregulation can reduce noise in the expression of a transcriptional repressor, relatively little attention has been paid to whether or not this is an effective strategy for reducing noise in downstream genes regulated by the same repressor. Such a network was found to reduce noise in *Saccharomyces cerevisiae*^9^, but in *Escherichia coli* repressor expression noise could be relatively high (e.g. from relatively small cell volume, short mRNA lifetime, and high extrinsic noise), and amplified downstream in transcriptional cascades^12-14^. To ensure that noise reduction in repressor expression propagates to the gene of interest, one could express it in fusion with a repressor and proteolytically cleave to achieve one-to-one expression^10^. Alternatively, bicistronic expression can allow for different expression levels of repressor and the gene of interest while eliminating transcriptional noise. Polycistronic transcription is a common motif in operons shown to reduce noise in genetic networks^15,16^ and to be especially important in efficient production of heteromeric protein complexes^17,18^.

Autoregulatory, bicistronic expression systems have been implemented in cell-free expression systems^19^ and shown to partially compensate for plasmid copy number variation in *Escherichia coli*^6^. However, gene expression noise for such systems has not been analyzed experimentally and, despite the apparent potential for reducing noise in the expression of a gene of interest, autoregulatory, bicistronic gene expression is not commonly used to control recombinant protein expression. I will show with stochastic simulations using parameters typical for *Escherichia coli* gene regulation that such an expression system produces a relatively linearized inducer dose-response, with noise below the “extrinsic noise limit” observed for chromosomal *Escherichia coli* genes^1^. I will then introduce one such system implemented on a plasmid and characterized its dose-response and noise level, showing that ribosome binding site modification can predictably expand the available dynamic range. Experimental comparison to alternative regulatory circuits confirms that bicistronic autoregulation reduces gene expression noise. Finally, I propose a hybrid system that reduces noise to the extrinsic noise limit while greatly expanding the available dynamic range.

## Methods

### Stochastic simulations

Custom MATLAB (The MathWorks, Inc.) scripts were made to simulate exact trajectories of the reactions in Table 1 using the Gillespie next-reaction stochastic simulation algorithm^20^ with unit volume. Note that with unit volume, concentrations and molecule numbers are equivalent, and all reactions have units of s^−1^. An exact algorithm is important as many species are at low numbers. All reactions were included in all simulations, and the configurations shown in Fig 1A could be realized within the same framework by setting rates of some reactions to zero (Table 1 indicates the reactions included for each configuration). Table 1 lists reaction rates used. The bimolecular association rate of 10^−5^ s^−1^ is within the range of those used for typical microbial cell volumes on the order of 1 ~m^3^. Transcription rate *k*_TX1_ was set to 1.7x10^−3^ s^−1^ for all simulations except for when it was lowered to simulated weakened, constitutive expression by lowering *k*_TX1_ to 6.5x10^−6^ s^−1^. Transcription rate *k*_TX2_ was also set to 1.7x10^−3^ s^−1^ except for in the hybrid repressor scheme where it was 6.7x10^−4^ s^−1^. Degradation rates for mRNA and protein were chosen to match typical *Escherichia coli* mRNA lifetimes cell growth rates, respectively. Degradation of inducer-bound repressor results in liberation of one inducer molecule; DNA-bound repressor is protected from degradation. The translation rate *k*_TL_ was set to 0.67 s^−1^ except when reduced to simulate weakened ribosome binding sites.

**Table 1.**
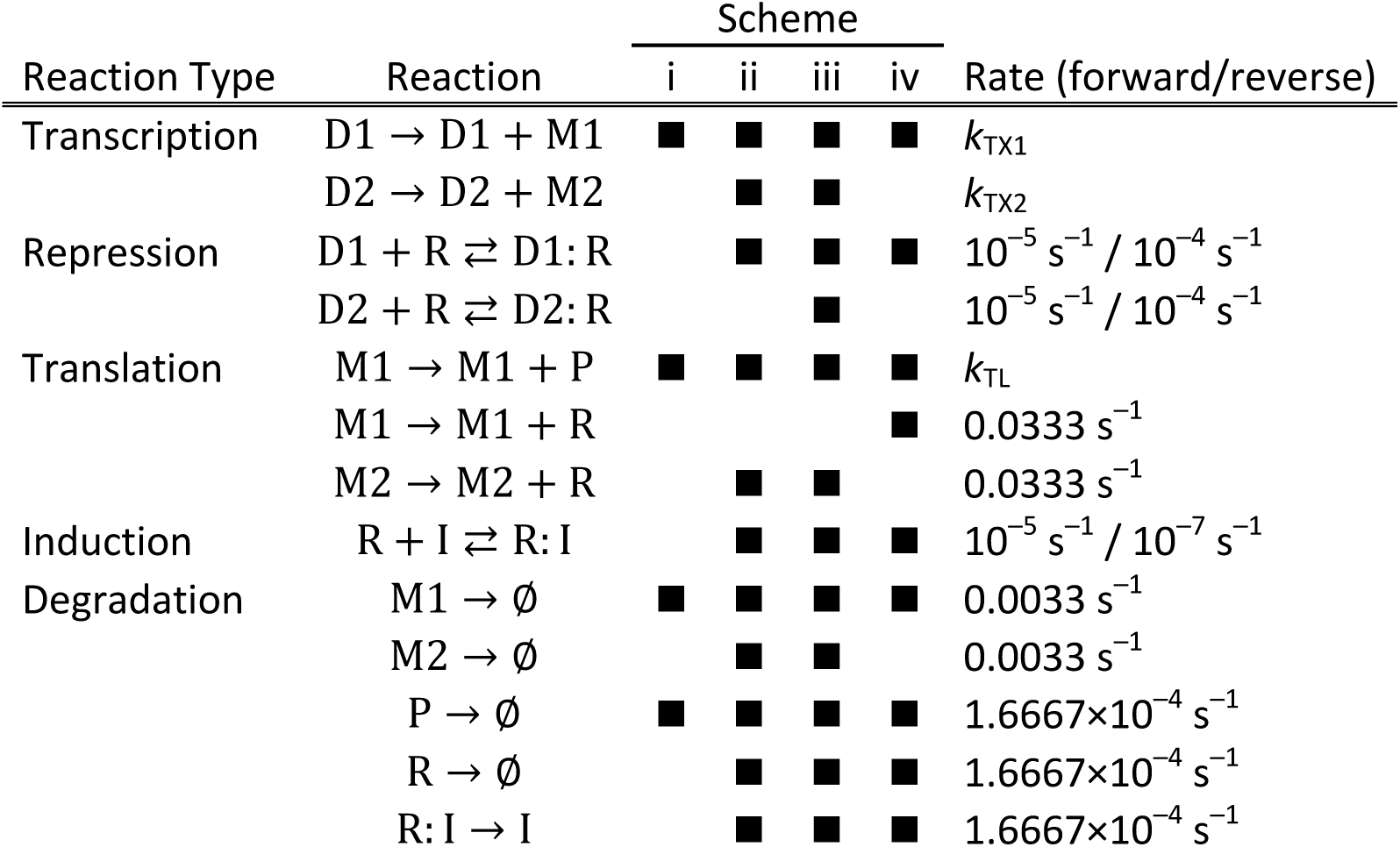
List of chemical reactions in stochastic simulations and reaction rates. The network schemes in Fig 1A can be simulated by including the reactions with black squares for (i) constitutive expression, (ii) constitutive repressor, (iii) autoregulated repressor and (iv) bicistronic autoregulation. Rates fixed in all simulations (when not set to zero) are listed below along with names of variable reaction rates. The number of free inducer molecules was kept constant. **S1 Text. Supporting Material.** DNA sequences for all constructs, Tables A and B, Figures A— F.

**Fig 1.**
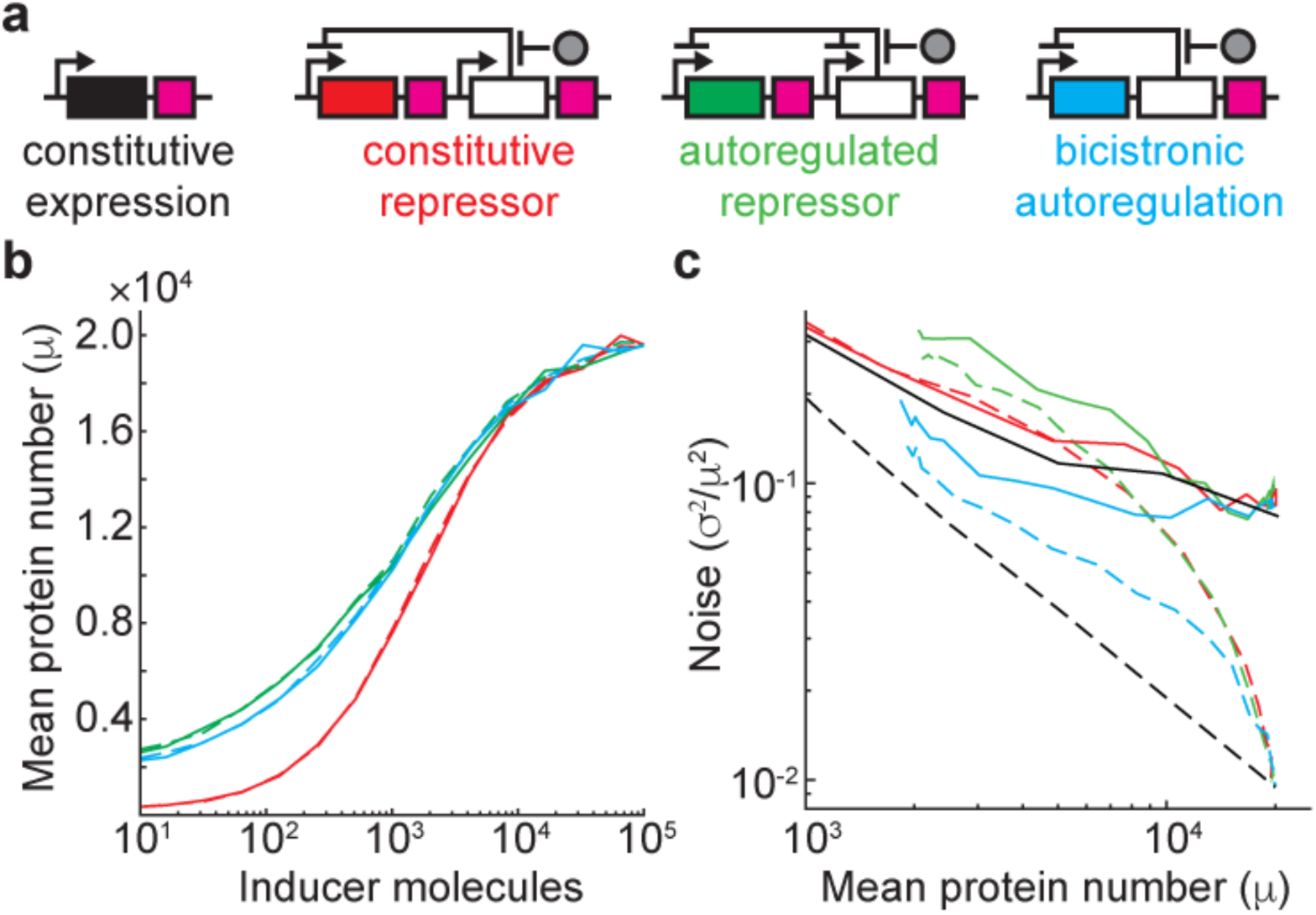
Simulated gene expression dose-response and noise in different regulatory circuits. Schematics of the regulatory schemes explored by stochastic simulation. Genes of interest (including coding sequence and translation start/stop signals) are shown as rectangles colored black (constitutive expression), red (repressed by a constitutively expressed transcriptional repressor), green (repressed by an autoregulated transcriptional repressor) and blue (autoregulated, bicistronic expression). White rectangle, transcriptional repressor. Gray circle, inducer. Black arrow, promoter. Purple square, transcription terminator. (**B**) Response to inducer for all regulated networks in the absence (dashed lines) and presence (solid lines) of extrinsic noise. Lines are colored according to the gene colors in Fig 1A. (**c**) Dependence of gene expression noise on average expression. Line style same as in Fig 1B.

Simulations started with 10 unrepressed DNA copies, specified numbers of inducer molecules, and no additional molecules. Inducer molecules were maintained at a constant free concentration, consistent with rapid equilibration of intracellular and extracellular volumes. Simulations at the limit of slow equilibration (repressor-bound inducer is never replaced) gave qualitatively similar results, with a larger noise reduction for the bicistronic, autoregulated system (**Fig A in S1 Text**). Simulation were run for 101,000 minutes with the system state stored every 1 minute. To implement extrinsic noise, an Ornstein-Uhlenbeck process with parameters τ=200 min, c=2.5x10^−5^ was generated^21^. Following previous work^22^, this time series was exponentiated and scaled by its mean. The resulting value was used to scale all translation rates. The first 1,000 minutes of all simulations were excluded from analysis to allow simulations to reach equilibrium, which occurs after a few hundred minutes (**Fig B in S1 Text**). 100,000 minutes of simulation time was chosen to balance computer time with acquiring continuous and reproducible noise measurements.

### Plasmid engineering

A plasmid was synthesized by Genewiz (New Jersey, USA) by inserting a synthetic sequence into the pUC57-Amp vector. The high-copy pUC origin of replication was replaced by p15a (estimated at ~18–30 copies per cell^23,24^). In the pZH501 plasmid, a non-fluorescent protein is expressed (a fusion of the bacteriophage lambda protein CI and SNAP-tag^25^). The CI-SNAP ORF in this plasmid was replaced by GFPmut2^26^ to create pZH509; GFPmut2 is referred to as “GFP” throughout this manuscript. The full DNA sequence of region encoding inducible GFP expression is included in S1 Text. It includes the hybrid P_LtetO-1_ promoter (containing bacteriophage λ P_L_ promoter overlapped by two copies of the tetO_2_ sequence)^23^, open reading frames with independent ribosome binding sites and double stop codons for GFP and *tn10* TetR^27^, and the *rrn*B T1 transcription terminator^28^. Weakened ribosome binding sites were designed using an online ribosome binding site calculator^29^ to generate plasmids pZH510, pZH511, pZH512 and pZH513 (**Table A in S1 Text**). Estimated RBS strengths were calculated using the reverse engineering mode of the RBS Calculator (v2.0, available at http://www.denovodna.com) using the first 839 bp of the mRNA sequence transcribed from P_LtetO-1_ including the GFP CDS and the first 50 bp of the TetR CDS. The predicted RBS strength for TetR was 867, so the number of TetR molecules per cell is predicted to range from 6.7% (pZH509) to 59% (pZH511) of the number of molecules of GFP (taking into account only the predicted RBS efficiencies for each CDS). Plasmid modification was done using PCR and 1-or 2-fragment isothermal assembly^30^. Plasmids were transformed into *Escherichia coli* TOP10 (Invitrogen) for fluorescence experiments.

Constitutive GFP expression plasmids pZH514, pZH515 and pZH516 were generated from inverse PCR of templates pZH509, pZH511 and pZH512, respectively, to eliminate TetR. Monocistronic, autoregulated plasmids pZH517, pZH518 and pZH519 were generated by isothermal assembly of one fragment amplified by inverse PCR of pZH509, pZH511 and pZH512, respectively, and another fragment containing the *rrnB* T1 terminator and the P_LtetO-1_ promoter. Plasmids pZH520, pZH521 and pZH522 with constitutively expressed TetR were generated following the same protocol except the inserted contained the moderate-strength, constitutive proB promoter^31^. Insert DNAs were synthesized by IDT (Iowa, USA). All plasmids were verified by sequencing and sequences are available in S1 Text.

### Growth conditions

For most experiments, cells were grown in 1 mL cultures of M9A minimal media (48 mM Na_2_PO_4_, 22 mM KH_2_PO_4_, 8.6 mM NaCl, 19 mM NH_4_Cl, 2 mM MgSO_4_,0.1 mM CaCl_2_) supplemented with 50 μg/ml carbenicillin, 1X MEM amino acids (Without L-Glutamine, Life Technologies 11130-051) and 0.4% glucose (“M9A” medium) at 30°C with shaking in 14-mL polypropylene culture tubes. Overnight cultures were diluted to OD600=0.02 and maintained in exponential growth until observation by flow cytometry or microscopy. For steady-state induction experiments cells were grown at least 4 h in the presence of inducer before observation. TetR and GFP expression was induced by the addition of anhydrotetracycline hydrochloride (ATc, diluted from 100 ~M stock in 50% ethanol). When necessitated by long experiments (e.g. timelapse flow cytometry), cell cultures were occasionally diluted in fresh media kept at 30°C to maintain exponential growth. For flow cytometry experiments using the BioRad S3e cell sorter, cells were grown in M9A media supplemented with 1% rich SOB media (2% tryptone, 0.5% yeast extract, 8.6 mM NaCl, 2.5 mM KCl, 10 mM MgCl_2_) and 50 μg/ml carbenicillin at 37°C.

### Flow cytometry (BD Accuri C6)

Flow cytometry experiments utilized the BD Accuri C6 sampler equipped with a 24-tube robotic sampler; this device has a linear response to fluorescence intensity and is regularly calibrated using standardized fluorescent beads. Samples were harvested during exponential growth (OD600=0.1—0.3), diluted between 1:33 and 1:100 in 1 mL PBS pH 7.4 depending on culture density, and sampled using the “fast” flow setting to capture 100,000 events above a forward scattering height threshold of 7,000. Samples were agitated between each collection time. The sum of the autofluorescent background and GFP expression was taken to be proportional to peak area of the FL1 detector using 488-nm laser excitation and a 533/30 nm emission bandpass filter.

For induction and repression kinetics experiments, ATc was added to uninduced TOP10/pZH509 cells to 4 nM and samples were taken every ten minutes, increasing ATc to 8 nM and then to 16 nM after 90 and 170 minutes, respectively. The 16-nM ATc sample was then centrifuged and washed twice in media lacking ATc before taking samples every 10 minutes to observer repression kinetics; there was a 10-minute delay between beginning the wash protocol and acquiring the first flow cytometry sample.

For all flow cytometry experiments, events were gated after data acquisition to fall near the peak in the histogram for forward-scattering area (FSCA) and side-scattering height (SSCH) (**Fig C in S1 Text**). FSCA and SSCH were empirically found to maximize the ability to discriminate between weakly scattering *Escherichia coli* cells and background events. For all samples approximately one third of events satisfied the criteria:

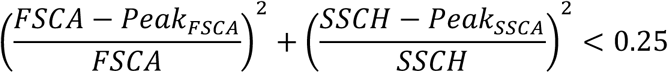

All analysis shown in Fig 2 used the mean fluorescence intensity from this gated sample, without subtracting the autofluorescence background of 191 (**Fig D in S1 Text**). When comparing ribosome binding sites, fluorescence intensity distributions were fit as the convolution of the autofluorescence distribution (from non-fluorescent strain pZH501) and a log-normal distribution (**Fig D in S1 Text**). The reported GFP expression mean is the mean of the fit log-normal distributions.

**Fig 2.**
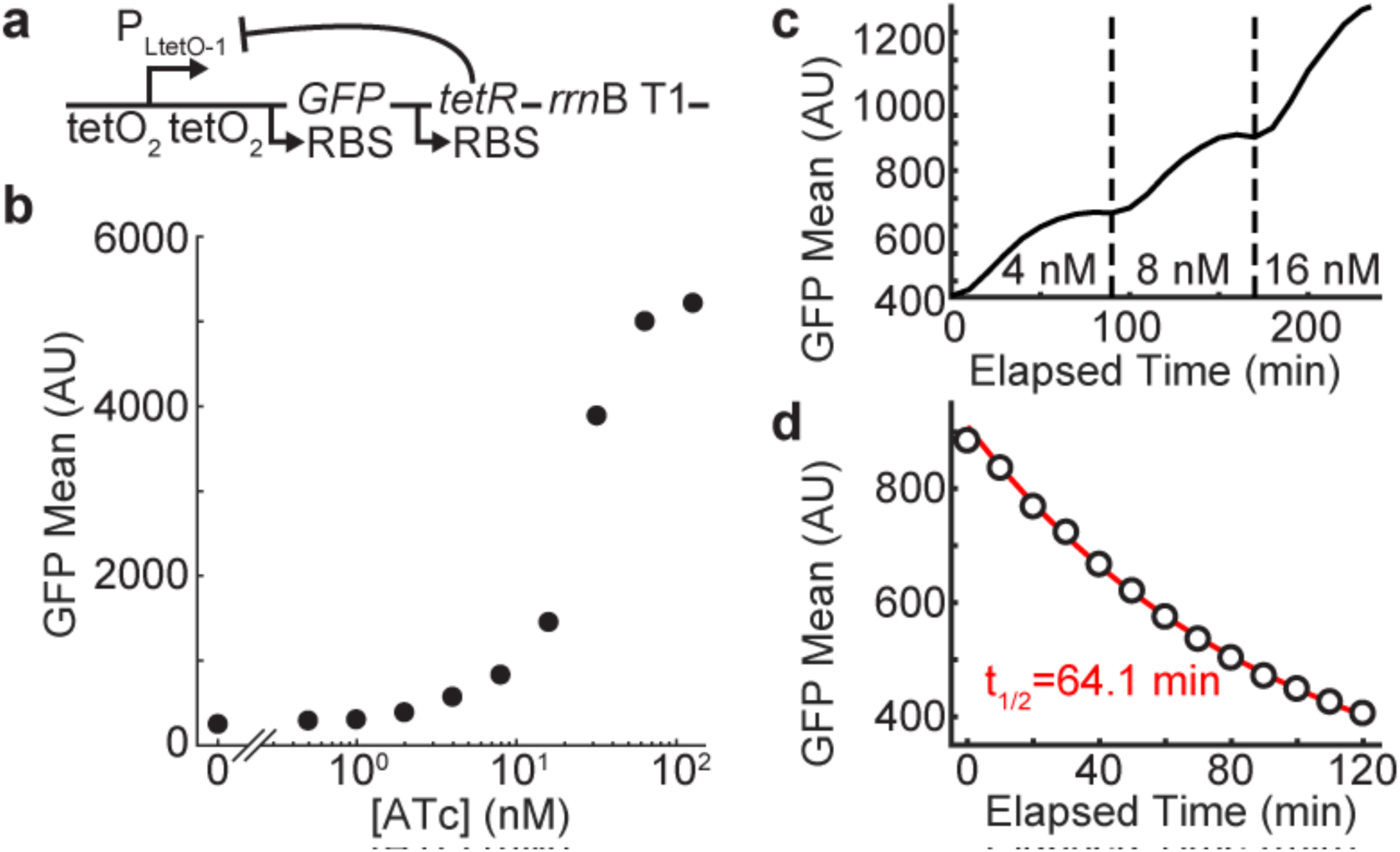
Characterization of induction dose-response and dynamics for bicistronic autoregulation circuit. Schematic of autoregulatory construct, carried on a plasmid. The P_LtetO-1_ promoter encodes the bicistronic transcript for GFP and TetR and is terminated at *rrnB* T1. TetR binding to either of two tetO_2_ sites represses transcription. (**B**) Flow cytometry analysis of pZH509 induction. Approximately a 50-fold change in expression level across induction range (0—128 nM ATc), with an 8-fold change in ATc (8 to 64 nM) giving a 6-fold increase in GFP expression. (**C**) Induction kinetics for pZH509 measured by flow cytometry; mean GFP fluorescence with ATc increased from 0 to 4 nM at 0 min, 4 to 8 nM at 90 min, 8 to 16 nM at 170 min In all cases the half-time of approaching equilibrium is ~30 min (occurring at ~30, 120, and 200 min). (**D**) Repression kinetics. Cells grown in 16 nM ATc were washed starting at at t = –10 min and observed starting at t = 0 min. The decrease in fluorescence from t = 10 min to t = 120 min is well fit by exponential decay with a half-time of 64.1 min.

### Flow Cytometry (BioRad S3e)

Noise comparison to alternative regulatory constructs was measured using the BioRad S3e cell sorter to assay cellular GFP fluorescence (488 nm excitation with 525/30 nm bandpass filtered emission). This device has a linear response to fluorescence intensity and is calibrated daily using fluorescent beads. Overnight cultures were diluted 1:100 and grown for 2.5 hours at 0, 1, 2, 4, 8, 16, 24, 32, 40, 64, 128 and 256 nM ATc (the negative control, pZH501, and the constitutive expression strain pZH514 was grown at 32 nM aTc). Samples were collected at a target of 2,000 events per second for 30,000 events (FSC gain 400, threshold 0.5; SSC gain 280; FL1 gain 650). Data was gated according to the procedure above, except that a FSCA/SSCH threshold of 0.5625 rather than 0.25 was used to gate approximately one third of events. Noise and mean were estimated directly from the integrated GFP intensity (FL1A) of events. All samples except for 0 nM ATc, pZH509 were well above background so it was unnecessary to account for autofluorescence.

### Microscopy

Cells growing exponentially were spotted onto M9A agarose gel pads (3% SeaPlaque GTG, Lonza) also containing ATc and imaged on a Elyra PS1 microscope (Zeiss) using a 100 mW, 488 nm excitation laser, 100x/1.46 a-plan apochromat oil immersion objective, and Andor Ixon DU 897 EMCCD. To quantify the intensity of single GFP molecules, cells were imaged continuously at 100% laser power with 17.5 ms integration times in a 128x128 pixel^2^ area. Although most GFP molecules were rapidly diffusing in the cytoplasm, a sufficient number of molecules produced diffraction limited spots that were detected and fit to a Gaussian PSF using Fiji^32^ with the ImageJ^33^ plug-in Thunderstorm^34^ (v1.3) with standard parameters for wavelet (B-spline) decomposition with a 4-pixel fitting radius and a detection threshold of “std(Wave.F1)*1.5”. Integrated fluorescence intensities for 858 detected spots were fit to a gamma distribution giving a mean spot intensity of 3,669 counts (**Fig E in S1 Text**). After verifying that *in vivo* fluorescence intensity was linear with respect to integration time and laser intensity, this was scaled to intensities of 1,101 and 110.1 counts for 35-ms images at 15% and 1.5% laser power, respectively.

Dark background was subtracted from fluorescence images and images were flattened to adjust for uneven illumination following a previously reported procedure^1^. Cells were manually segmented from brightfield images in Fiji^32^ and the integrated fluorescence was calculated from the same region in the fluorescence image before being divided by number of counts per single GFP molecule to estimate the total number of molecules per cell. To normalize for cell size to allow comparison with a previous measurement of extrinsic noise^1^, the number of molecules per cell was divided by the cell area and then multiplied by the mean cell area (area in brightfield images is approximately proportional to cell volume for cylindrical *Escherichia coli* cells). Following the procedure for flow cytometry, data was also collected for the non-fluorescent pZH501 strain, and distribution statistics were estimated by fitting the convolution of the background fluorescence histogram and a log-normal distribution. Error in log-normal fitting was assessed by fitting 100 equally sized data sets generated by bootstrap sampling with replacement (**Table B in S1 Text**).

## Results

### Simulating inducer dose-response and gene expression noise

Four gene regulatory circuits (Fig 1A) were modeled in stochastic simulations incorporating transcription, translation, repressor binding, inducer binding and mRNA/protein degradation. Reaction rates were chosen to give protein and mRNA numbers and lifetimes typical for prokaryotic cells, with maximal repressor expression levels similar to those reported (1,000 per cell) for typical inducible gene expression systems^23^. Repression was modeled as a monomer binding DNA to inhibit transcription, with the binding of one inducer molecule to the repressor preventing DNA binding, and all simulations included 10 DNA copies to mimic plasmid-based gene expression. Extrinsic noise was simulated as a stochastic variation in translation rate(s)^22^. The four modeled regulatory circuits include constitutive expression, repression by a constitutively expressed transcriptional repressor (a common system used for inducible gene expression), repression by an autoregulated transcriptional repressor, and autoregulated, bicistronic expression of a gene of interest and transcriptional repressor.

One advantage of autoregulated gene expression is a less steep inducer dose-response (i.e. a “linearized” dose response), which can make it easier to achieve intermediate expression levels between maximal repression and maximal induction. Gene expression controlled by autoregulated TetR has been observed experimentally in yeast, *Escherichia coli*, and cell-free systems to have a linearized inducer dose-response relative to systems with constitutively expressed TetR^7,9,19,23^. Fig 1B shows the inducer dose-response for simulated, inducible systems. Indeed, adding negative feedback expands the inducer dose-response by about 1 order of magnitude of inducer concentration. The bicistronic system exhibits slightly lower expression at low-to-intermediate inducer concentrations, but otherwise exhibits a similar inducer dose-response monocistronic system. Inducer dose-response is unaffected by extrinsic noise in translation rate.

In Fig 1C, the regulated systems are compared to unregulated expression in which the transcription rate is varied to adjust mean expression levels. Protein noise in this simple, unregulated network depends on the number of proteins produced per transcript^8^, *b*, which is the same for all simulations. The dashed black line for unregulated expression without extrinsic noise is the intrinsic noise limit^8^, 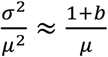 In the absence of extrinsic noise, all regulated schemes exhibit noise above the intrinsic noise limit. However, bicistronic expression is required to achieve noise levels near the limit, while an autoregulated repressor that separately represses the gene of interest exhibits as much or more noise than with a constitutively expressed repressor.

Extrinsic noise leads to a plateau in gene expression noise at high expression levels^10^. This is evident in Fig 1C with all simulations including extrinsic noise converging on the same noise value. A proteomic study in *Escherichia coli* found that gene expression noise converges to a global extrinsic noise limit of ~0.1^1^. Again, the constitutive repressor and autoregulated repressor simulations exhibit similar noise levels above the extrinsic noise limit; even though autoregulation can combat extrinsic noise^10^, this noise reduction is not transmitted to the downstream gene. However, implementing bicistronic expression again reduces noise—this time below the extrinsic noise limit. In the limit of slowly equilibrating inducer (inducer is not replenished from the environment after binding repressor), there exists a U-shaped inducer response that has been observed experimentally for autoregulated gene expression (**Fig A in S1 Text**)^7^. These simulations omit many molecular details of gene regulation and cellular heterogeneity that influence gene expression noise, so the principles of bicistronic autoregulated gene expression were implemented experimentally to see if noise could be reduced below the extrinsic limit.

### Construction and characterization of an autoregulatory, bicistronic gene expression system

A schematic of the bicistronic, autoregulated system is shown in Fig 2A in which TetR and GFP are bicistronically expressed from a TetR-repressible promoter. It is harbored on a plasmid with the p15a origin of replication, and it is similar in many aspects to systems previously used in *Escherichia coli*, *Saccharomyces cerevisiae*, and cell-free gene expression^7,9,19,23^. GFP expression during exponential growth (doubling time 63 minutes grown in M9A medium at 30°C) was monitored by flow cytometry and fluorescence microscopy. Here, expression was induced by anhydrotetracycline (ATc), a tetracycline analog that binds TetR more tightly^35^, in order to avoid tetracycline toxicity. However,tetracycline or other weaker-binding analogs may be preferable in order to minimize the effect of noise in intracellular inducer concentrations.

The inducer dose-response (Fig 2B) for the system was measured by flow cytometry. It is similar to that observed for autoregulated TetR expression in *Escherichia coli*^7^, with a response from ~1—100 nM ATc of almost two orders of magnitude of protein expression. This response is linearized compared to that of the same promoter regulated by constitutively expressed TetR^23^, where a 5-fold increase in ATc gave over a 5000-fold change in protein expression. Induction and repression kinetics were also characterized by flow cytometry. Equilibrium is re-established approximately 30 minutes after stepwise increases in ATc concentration (0, 4, 8, 16 nM), showing this system could be useful for experiments requiring dynamic transgene expression (Fig 2C). Cultures with 16 nM ATc were washed and GFP fluorescence decayed with a half-time of 64.1 min (Fig 2D). Reduction in GFP at the nearly same rate as cell growth indicates that repression is quickly re-established after ATc removal. These results confirm the utility of the bicistronic autoregulatory construct for precisely expressing a transgene at a desired level and rapidly changing induction levels.However, this comes at the cost of greatly reduced dynamic range.

### Expanding dynamic range through ribosome binding site modification

In order to expand the dynamic range of the bicistronic autoregulated expression system to cover the useful range for *Escherichia coli* transgene expression (~10—10,000 molecules/cell), weakened ribosome binding sites were designed^29^ for GFP in order to have a lower relative level of GFP expression versus TetR from the bicistronic transcript. Simulation results in Fig 3A show the predicted results of weakening ribosome binding sites by reducing translation rates. In the absence of extrinsic noise, noise as a function of mean expression decreases because of the reduced number of proteins produced per mRNA. In the presence of extrinsic noise, noise is observed to go below the extrinsic noise limit (noise at fully induced expression level) at intermediate inducer concentrations for all translation rates.

**Fig 3.**
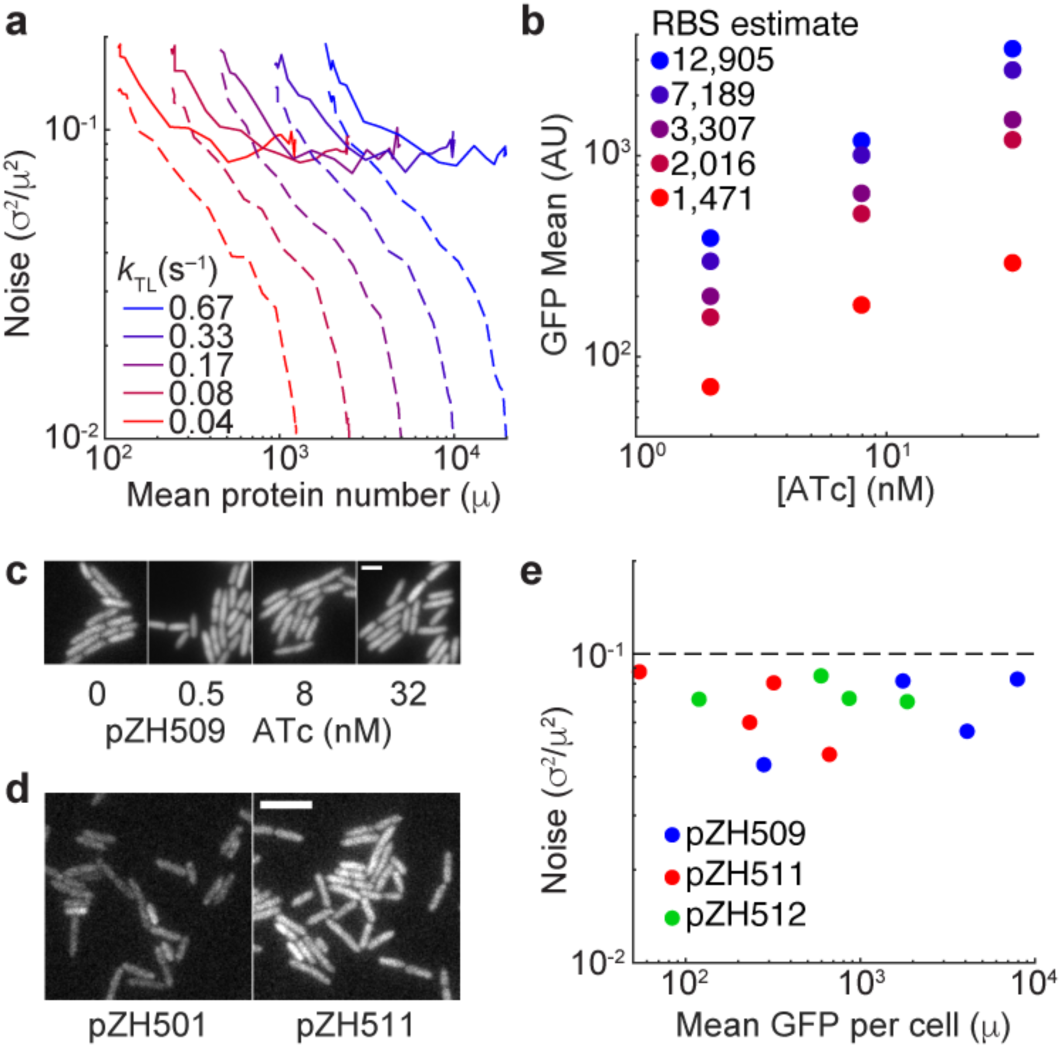
Induction dose-response and noise characterization for bicistronic autoregulation circuits with a range of expression levels. (**A**) Simulated gene expression mean and noise for the bicistronic, autoregulatory construct with different translation rates in the absence (dashed lines) and presence (solid lines) of extrinsic noise. (**B**) GFP expression means for strains with mutated GFP ribosome binding sites at various 2, 8, and 32 nM ATc. Strains are colored by predicted ribosome binding site efficiencies. (**C**) GFP expression in unrelated cells for the highest-expression strain pZH509 at 0, 0.5, 8 and 32 nM ATc. Maximum fluorescence intensities normalized by the mean number of GFP molecules per cell (**Table B in S1 Text**). Scale bar 2 ~m. (**D**) Fluorescence of the non-GFP-expressing plasmid pZH501 is compared to the lowest-expression-level plasmid pZH511 without induction. Intensity scaling identical for both images. Scale bar 5 ~m. (**E**) GFP expression mean (molecules/cell normalized by cell area) and noise for plasmids pZH509, pZH511, and pZH512 at 0, 0.5, 8 and 32 nM ATc. Dashed line indicates the approximate global extrinsic noise limit^1^.

This strategy was implemented by designing 4 new ribosome binding sites for a total of 5 different strains. Fig 3B shows expression levels at 3 different ATc concentrations compared to the predicted translation efficiencies, which are expected to scale linearly with protein expression level^29^. Experimental expression levels were observed to monotonically increase with predicted translation efficiencies (Table 1 **in S1 Text**). Three plasmids with different expression levels were chosen to use to measure GFP expression noise and expression levels by fluorescence microscopy: the original construct pZH509, plus pZH511 and pZH512 with expression levels of 15% and 43% those of pZH509, respectively.

### Reducing gene expression noise below the extrinsic noise limit

GFP expression noise in pZH509, pZH511 and pZH512 was quantified by fluorescence microscopy calibrated by single-molecule GFP intensities following a protocol previously applied to the *Escherichia coli* proteome^1^. Mean GFP expression and expression noise were estimated by fitting the observed fluorescent signal to the convolution of a log-normal distribution of single-cell GFP expression and autofluoresence of a negative control. Comparing unrelated cells spotted onto an agarose gel pad, it was immediately apparent that GFP expression noise was much lower than that typical for constitutive expression from a plasmid at a range of ATc concentrations for the high-expression pZH509 strain (Fig 3C, 283—7,990 molecules/cell).Low expression noise was evident even for the lowest expression level where autofluorescence significantly contributes to the total signal (Fig 3D, 55 molecules/cell).

Gene expression noise was estimated from the distributions of GFP molecules per single cell, normalized by cell size (Fig 3E). Remarkably, noise was below the previously observed extrinsic noise limit for chromosomal genes of ~0.1^1^, across the expression range investigated (55—7,990 molecules/cell). This is despite the fact that gene-dosage noise is expected to be greater for expression from a plasmid than from the chromosome.Combining mean expression level measurements from flow cytometry and microscopy expression levels, a range of 55 (pZH511, 0 nM ATc) to 10,740 (pZH509, 128 nM ATc) is achievable. It is outside the scope of this manuscript, but it is expected that an increase in noise will be observed at high ATc concentrations; where there is effectively no autoregulation, noise will increase to the extrinsic limit. It is possible to design a stronger ribosome binding site than that in pZH509, but expression levels of a single gene beyond~10,000 molecules/cell could perturb behavior. It is also possible to design a weaker ribosome binding site than that in pZH511, but in that case it is very important to control for possible alternative, in-frame ribosome binding sites.

Although this experiment shows lower noise than the previous measurement of ~0.1, differences between cell growth protocols and fluorescent proteins could affect measured gene expression noise. To address this concern, I created plasmids with the alternative regulatory constructs shown Fig 1A. Constitutive expression was observed from the strong, unrepressed P_LtetO-1_ promoter; other constructs were observed at a range of induction levels. I compared GFP induction (Fig 4A) and found that the inducer dose-response was linearized in both strains with autoregulated TetR expression. Notably, the bicistronic autoregulation circuit exhibited lower maximal expression than other strains (**Fig F in S1 Text**), possibly indicating lower stability or a lower translation initiation rate for the polycistronic mRNA. Noise for the bicistronic autoregulation circuit was lower than that of the circuit in which GFP and TetR are transcribed separately, and below the extrinsic noise limit for strong, constitutive expression (Fig 4B). Unexpectedly, noise in the circuit with constitutively expressed TetR was very high at intermediate ATc concentrations, reflecting a bimodal distribution in GFP expression (**Fig F in S1 Text**). This deviation from the simulated results (Fig 1) possibly results from slow kinetics of ATc/TetR and TetR/DNA association as well as not accounting for cooperativity in the simplified network that was simulated.

**Fig 4.**
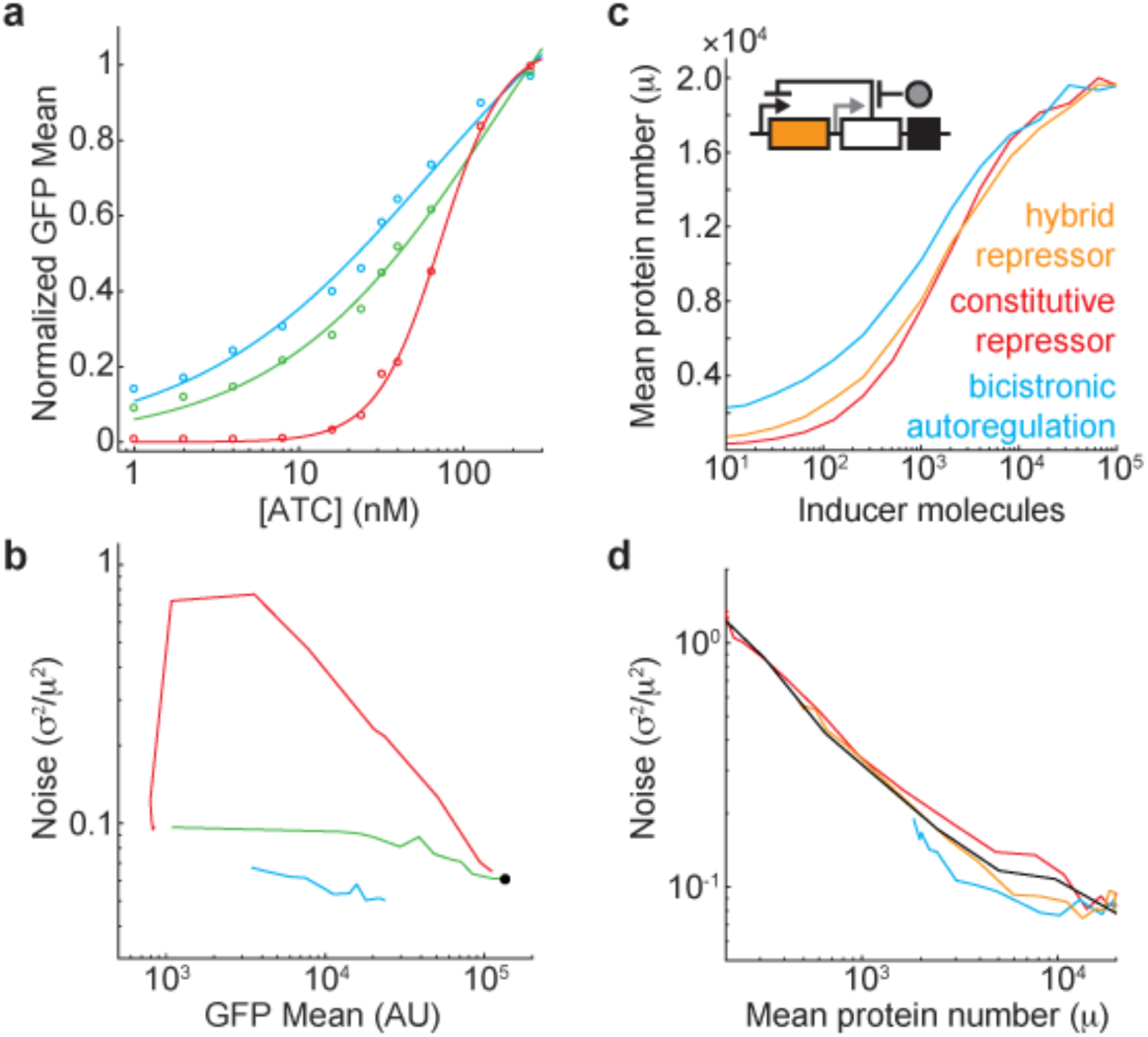
Comparison of regulatory circuits and increased dynamic range with a hybrid circuit. (**A**) GFP induction is measured by flow cytometry and fit using the Hill equation for pZH509 (blue, *n*_*h*_ = 0.60 +/-0.16), pZH517 (green, *n*_*h*_ = 0.65 +/-0.14) and pZH520 (red, *n*_*h*_ = 2.24 +/- 0.22). Data and fit curves are normalized to the fit value at 256 nM ATc. Data not shown for 0 nM ATc (**Fig F in S1 Text**). (**B**) Noise dependence on mean expression level; coloring identical to Fig 4A. Black dot, pZH514 at 32 nM ATc. Noise cannot be calculated for pZH509 at 0 nM ATc because of low expression (**Fig F in S1 Text**). (**C**) A hybrid scheme is proposed (inset) in which repressor (white box) expression occurs both from autoregulated (black arrow) and relatively weak (gray arrow) promoters that share a transcription terminator (black box). This achieves an inducer dose-response in the gene of interested (orange) that is less steep than in the absence of autoregulation (red) while increasing the dynamic range relative to bicistronic autoregulatory circuit (blue). (**D**) The hybrid system reduces noise relative to the system with constitutively expressed repressor, with noise at or below the extrinsic limit (black). All simulations include extrinsic noise.

## Discussion

The bicistronic, autoregulatory expression system can be easily adopted to many experiments and implemented in other organisms; replacing GFP with a gene of interest using modern polymerases requires two PCR reactions and one isothermal assembly step requiring a few hours and having nearly 100% efficiency. It is trivial to construct orthogonal systems using alternative transcriptional repressors for expressing multiple genes. Care must be taken to calibrate expression levels in all experiments: one cannot simply replace GFP in one of these plasmids with a gene of interest and assume similar expression rates, because translation efficiency of both the gene of interest and TetR is dependent upon sequences near the ribosome binding sites. However, these concerns apply to all recombinant gene expression systems.

The largest noise reduction relative to unregulated expression (Fig 1C) occurs at intermediate induction levels. Thus, to minimize noise one should choose a ribosome binding site that gives desired expression level within the intermediate induction range (~20—30 nM ATc). This contrasts with systems regulated by constitutively expressed TetR, where bimodal expression at intermediate ATc concentrations can lead to a peak in gene expression noise^7,9^. It is striking that one of the most common methods for inducing a targeted level of recombinant gene expression—induction by inhibition of a constitutively expressed transcriptional repressor—utilizes a motif proven to increase gene expression noise. Despite the potential for an underlying bimodal gene-expression distribution, countless experiments using this induction method employ population averaged assays to measure protein expression levels that are then applied to cellular-scale models.Autoregulated, bicistronic expression avoids this problem. Here, I implement this bicistronic, autoregulation approach in *Eschericia coli*, where polycistronic expression is common. In principle, it can also be implemented in other organisms using internal ribosome entry sites (IRES) or by inserting small, self-cleaving amino acid sequences between the induced gene and the transcriptional repressor^36^.

I have shown that negative autoregulation is valuable for linearizing the inducer dose-response and reducing gene expression noise. However, this comes at the expense of dramatically reduced dynamic range (compare over 5,000-fold for the P_LtetO-1_ promoter repressed by constitutively expressed TetR^23^ to approximately 50-fold for pZH509, with similar results in the simulations in Fig 2B). In Fig 4C and Fig 4D I suggest a hybrid gene expression system in which a repressor is transcribed both constitutively and from a bicistronic, autoregulated promoter along with the gene of interest. The behavior of the system is expected to transition from that of the bicistronic, autoregulated system (with an infinitely weak constitutive promoter) to that of a system with a constitutively expressed repressor (with a relatively strong constitutive promoter). Simulations with the addition of a constitutive promoter that has a transcription rate 40% that of the bicistronic promoter show that this system recovers much of the lost dynamic range while still exhibiting reduced noise at or below the extrinsic noise limit. Engineering such an expression system will benefit from the development of promoters insulated from surrounding sequences^31^ so thathe transcription rate from the constitutive promoter is minimally affected by either transcription from the inducible promoter or the upstream sequence. Such an expression system would be valuable in experiments requiring both low gene expression noise and a wide range of expression levels.

## Acknowledgements

This work was financially supported by: Project LISBOA-01-0145-FEDER-007660 (Microbiologia Molecular, Estrutural e Celular) funded by FEDER funds through COMPETE2020 - Programa Operacional Competitividade e Internacionalização (POCI), by national funds through FCT - Fundação para a Ciência e a Tecnologia, and by the Dean's Research Unit at the Okinawa Institute of Science and Technology (OIST). Experiments were done at OIST; data analysis, simulations and paper writing were done at the Instituto de Tecnologia Química e Biológica António Xavier. I thank Dr. Eder Zavala (University of Exeter) for his insightful comments on the manuscript.

## Data Availability

Raw microscopy and flow cytometry data (BD Accuri C6), MATLAB simulation code, and explanatory text files sufficient for reproducing the results in this paper are available at Zenodo (DOI: 10.5281/zenodo.569808). Additional flow cytometry data (BioRad S3e) is available at Zenodo (DOI: 10.5281/zenodo.944156). An annotated DNA sequence for the bicistronic, negative feedback construct is included in S1 Text.

